# Collagen Production and Niche Engineering: A Novel Strategy for Cancer Cells to Survive Acidosis and Evolve

**DOI:** 10.1101/711978

**Authors:** Mehdi Damaghi, Samantha Byrne, Liping Xu, Narges Tafreshi, Bin Fang, John M. Koomen, Aleksandra Karolak, Tingan Chen, Joseph Johnson, Nathan D. Gallant, Andriy Marusyk, Robert J. Gillies

## Abstract

Ductal Carcinoma in situ (DCIS) is an avascular disease characterized by profound acidosis. Pre-malignant cells within this niche must adapt to acidosis to survive and thrive. A component of this acid-adaptation involves extracellular matrix remodeling leading to niche construction and remodeling. Using discovery proteomics, we identified that collagen producing enzyme PLODs are upregulated in acid-adapted breast cancer cells. Second harmonic generation microscopy showed significant collagen deposition within DCIS lesions of patients. In vitro analyses identified that acid-adaptation involves production of rare collagens that can be regulated by Ras and SMAD pathway. Secretome analysis showed upregulation ECM remodeling enzymes such as TGM2 and LOXL2. Comparison of acid induced collagens *in vitro* and *in patient* data showed correlation between rare collagens production and survival of patients. We conclude acidosis induces collagen production by cancer cells and promote growth independent of basal membrane attachment. The independently produced collagen can be used for niche construction and engineering as an adaptation strategy of cancer cells to survive and evolve.

## Introduction

The earliest stages of carcinogenesis are not known with certainty. Breast cancers are initiated by intraductal hyperplasia which is likely triggered by chronic inflammation (1–5). This involves cells growing into the ductal lumen, which requires them to detach from their normal niche which is attached to a basement membrane (BM) (6, 7). The attachment of epithelial cells to the basement membrane substratum is necessary for their survival, which is promoted by collagen activation of focal adhesion kinases, FAKs (8, 9). The death caused by substrate detachment is termed *anoikis* and it is a pre-requisite for cells to lose sensitivity to anoikis in order to survive and grow away from the BM and towards ductal lumens (10, 11). Since the blood supply for these cells resides in the surrounding stroma, as hyperplasic cells grow to the lumen, they become increasingly hypoxic (12). This hypoxia, along with nutrient deprivation, selects for cells with the phenotype of aerobic glycolysis, also known as the Warburg Effect, WE (11, 13). A significant sequela of aerobic glycolysis is the overproduction of lactic acid (12, 14–16). Because intraductal neoplasias are avascular, this results in significant acidification of the extracellular microenvironment, accompanied by upregulation of transporters to export intracellular acids to the extracellular compartment (17–20). Acidosis, in normal cells, induces apoptotic cell death (21). In carcinogenesis, however, cancer cells must eventually adapt to this harsh acidic environment (4) that arises phenotypic resistance to acid-induced apoptosis (4, 22).

The structural component of tumor niche is mostly extracellular matrix (ECM). Most of what is known about the role of the ECM in tumor biology has come from studies of normal mammary gland development, or invasion of cancer cells into the surrounding stroma. Although ECM is primarily composed of water, proteins and polysaccharides, it shows exquisite tissue specificity as a result of its unique compositions and topographies e.g. tumor ECM is totally different than normal adjacent ECM (23). This unique structure of ECM is generated through a dynamic biochemical and biophysical interplay between the various cells in each tissue and their evolving microenvironment (24). Collagens are major proteins of the ECM. The unique mechanical properties of collagens are mainly controlled by its structure and density that affects ECM induced tumor growth, migration, and metastasis (25). Increased deposition and re-organization of collagens, fibronectin, and proteoglycans is observed in the transition from DCIS to locally invasive disease and also in later stages and metastasis (26, 27). It has been assumed that collagen in the ECM is derived from stroma cells such as fibroblasts and macrophages. Collagen surrounding normal ducts has a well-organized concentric structure. Tumor associated collagens are more heterogeneous, either being radially oriented towards vasculature or very unstructured, especially in hypoxic areas (28). Recent ideas indicate that collagen structures can provide a path for migration and invasion. Indeed, it has been shown that remodeled stiff collagens were used as invasion highways by glioma cells and breast cancer cells to migrate along the collagen fibers aligned towards the blood vessels (29, 30). ECM collagens can also be degraded by enzymes such as metalloproteinases (31) or remodeled through proteinase mediated processes by crosslinking enzymes such as lysyl oxidase, LOX (32) or protease independent pathways with, e.g. transglutaminase, TGM2 (33, 34).

In this work we identify an important component of niche construction and remodeling in early cancers: the acid-induced production of collagens by nascent pre-cancerous and locally invasive cancer cells. Herein, we identify by discovery proteomics, that adaptation of pre-malignant cancer cells to chronic acidosis involves upregulation of collagen producing (PLODs) and remodeling enzymes (LOXs and TGM2), which are required for collagen export from cells and which is a precursor to crosslinking with lysyl oxidase. Second Harmonic Microscopy (SHM) showed significant collagen deposition within the lumens of DCIS, which are devoid of stromal cells. RT-PCR showed that rare collagens are produced by acid-adapted pre-malignant and cancerous breast epithelial cells, and that these collagens provide a mechanism for resistance to anoikis. We also defined molecular mechanism of this adaptation and discovered how mutations such as Ras and its downstream pathway activation play role in defining the emerged phenotype. These discoveries will help us to understand the molecular mechanisms behind the emerging phenotypes in cancer and also how to design proper combination therapies in different disease stages and how different tumors respond in their evolutionary adaptation strategies, such as niche engineering and self-defined fitness, and the impact that these biological processes ultimately have on patient outcomes.

## Results

### Acid adapted cells are resistant to anoikis

It’s been known for more than a decade that the center of the ducts in solid cancer models such as breast or prostate cancer is one of the most acidic habitats in the whole tumor ecosystem (Fig 1A) (3, 4, 35). Mechanisms, allowing tumor cells to tolerate acidosis have been studied extensively by our group (4, 36, 37) and others (36, 38). However, the impact of acidosis on the microenvironment niche and its components such as collagens and cancer evolution are less understood. We have proposed that cell death imposed by loss of basement membrane attachment (“anoikis”) is one of the first evolutionary barriers that cancer cells reach in their early progression (19) and herein we investigated whether acidosis would effect this process. We frequently observe the acid adaptation marker (plasma membrane bound LAMP2b) overexpressed in the periluminal regions of early DCIS breast lesions (Fig 1A) prompting us to hypothesize that these acid-adapted cells are resistant to anoikis. To examine this question, we compared soft agar gel survival assay using parental MCF7 cells and their acid-adapted counterparts (AA MCF7) that have been grown in acidic (pH 6.5) media for more than 3 months; until their growth rate at this pH matches unselected cells at the physiological pH (pH 7.4). The results showed AA MCF7 cells grew significantly more and larger colonies in soft agar (Fig 1B) compared to non-adapted control cells (NA MCF7). To corroborate this finding, we examined anchorage independent growth of AA and NA MCF7 with the spherogenic assay in an ultra-low adhesion U shape plate. Whereas NA MCF7 cells only formed loose clumps containing relatively few cells; AA MCF7 formed large, well defined spheres (Fig 1C). The dramatic differences in colony shapes under low attachment cultures suggested that AA MCF7 have increased cell adhesion. These two experiments indicated that acid-adapted cells are capable of anchorage independent growth. To validate these observation, we analyzed previously generated SILAC proteomic data (Damaghi et al., 2015) from NA vs. AA MCF7 cells with GeneGo software; focusing on membrane proteins and extracellular matrix (ECM) components. We found that cell adhesion, ECM remodeling, and cytoskeleton remodeling proteins were elevated in AA MCF7 cells compared to non-adapted MCF7 (Fig 1D). Given the elevated expression of ECM components, as well as molecules responsible for cell-matrix adhesion, we asked whether AA MCF7 cells are capable of stronger adhesion to ECM using a spinning disk assay. To investigate this adhesive phenotype, we performed adhesion strength experiments on NA MCF7 and AA MCF7 cells as well as NA and AA MCF10A, and immortalized mammary epithelial cells, as a control. These experiments showed that AA MCF7 cells were more adhesive than NA MCF7 cells and this observation was reversed for AA MCF10A cells, implying differential matrix adhesion responses of cancer cells to acid adaptation, compared to normal cell adaptations (Fig 1E). Next, we asked whether increased adhesion to ECM, produced by AA MCF7 cells, might be responsible for an increased anchorage independent survival. First, we examined the activation status of focal adhesion kinase (FAK), a key signaling node modulating pro-survival effect of collagen adhesion. Indeed, we found that levels of activating phosphorylation of FAK were significantly higher in AA MCF7 cells (Figure 1F). Next, we asked whether an increased FAK activity was contributing to increased anchorage independent survival of AA MCF7 cells. To this end, we compared the impact of FAK inhibition on survival under 3D growth between AA and NA MCF7 cells. We found that FAK inhibitors PF-573228 and FAK Inhibitor-14 decreased the viability of AA MCF7 cells both in normal pH and acidic media, whereas NA MCF7 cells were unaffected, suggesting that increased cell adhesion to autonomously produced ECM might be responsible for increased anchorage independent survival during acid adaptation (Figure 1G).

**Figure 1:**
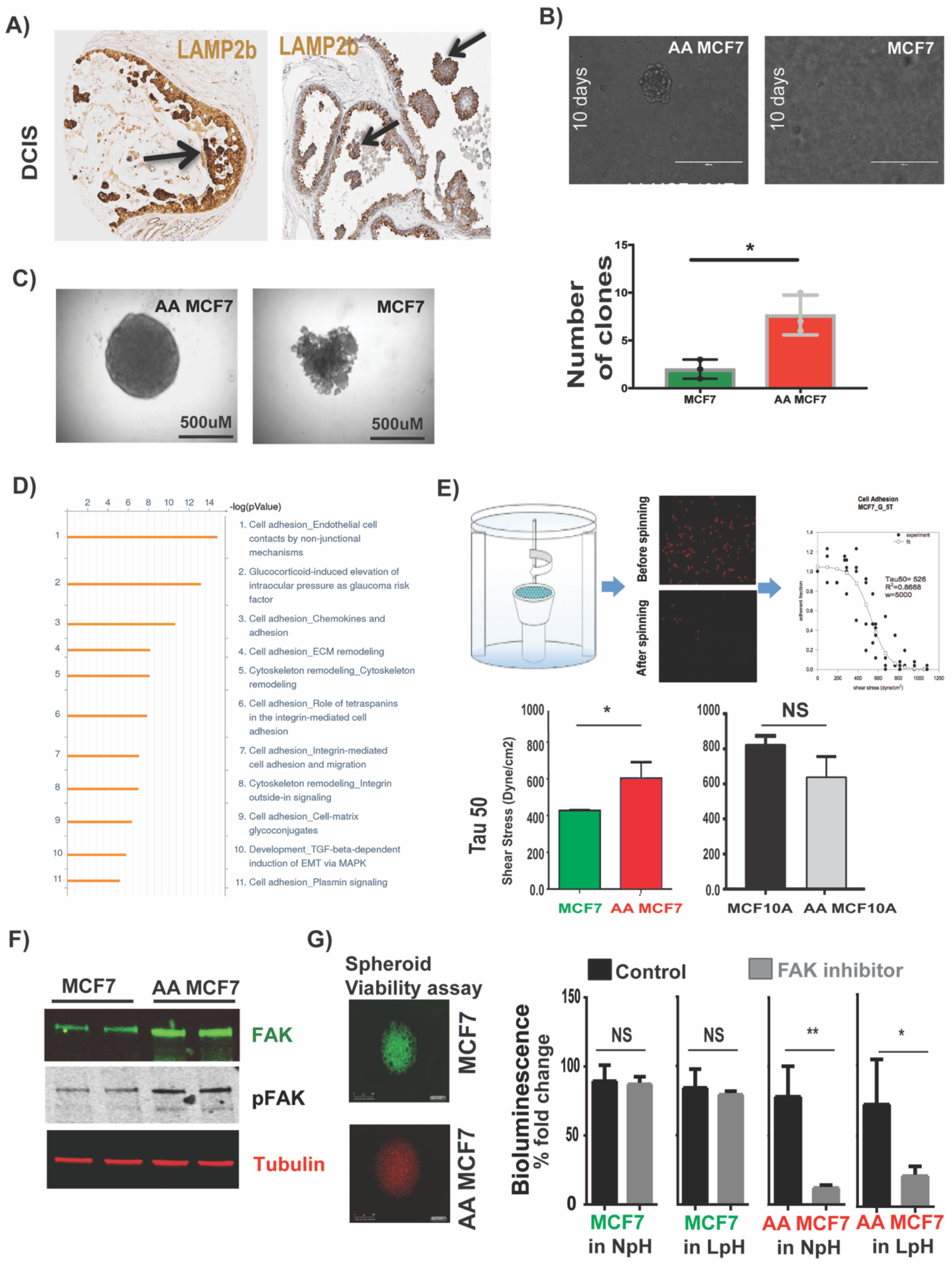
Acid adaptation prevents anoikis as a survival mechanism of early lesion cancer cells. A) Acid adapted cancer cells are frequently found in the center of the DCIS as one of the most acidic habitats inside the solid tumors. The staining is LAMP2b (acid adapted phenotype marker) on two DCIS patient biopsies. The highest expression of LAMP2b (the red color with mask) is located at the center of DCIS farthest from the vasculature. There are totally separated organoid inside the duct with no ECM contact at all with high amount of LAMP2b that is a marker of acid adapted cells. B) Soft Agar clonogenic assay on MCF7 and AA MCF7 cells. AA MCF7 cells grow more and bigger clones than non-adapted MCF7. C) Spherogenicty assay on MCF7 and AA MCF7 cells. Acid adapted MCF7 cells grow sphere in ultra-low adhesion plates contrary to the non-adapted cells. D) Proteomics analysis of AA MCF7 versus MCF7 cells. The analysis was applied on filtered proteins that have role in ECM and anoikis. More than 80 percent of the top ten pathways that were upregulated in acid adapted cells belong to cell adhesion, ECM-cell adhesion, and integrin mediated cell adhesion pathways implying the role of acid adaption in regulation of those pathways. E) Spinning disk experiment to measure cell-matrix adhesion. AA MCF7 cells are more adhesive than MCF7 while the normal MCF10A cells have opposite behavior. F) Western blot of FAK and pFAK on lystae from AAMCF7 and MCF7. G) 3D Viability assay of AA MCF7 and MCF7 spheroid treated with FAK inhibitors.

### Acid adaptation promotes collagen production of cancer cells

Above, we observed adhesion of cancer cells to matrix as a strategy to become anchorage independent for growth. However, within ductal lumens, it is expected that there would be no matrix for cells to adhere to as there are no matrix-producing fibroblasts. In ductal hyperplasia and carcinoma in situ, the only cells in peri-luminal volumes are expected to be epithelial-derived pre-cancerous and cancerous cells. Therefore, we hypothesized that acid-adapted cancer cells are either truly matrix independent, or they are producing their own matrix to allow survival. To probe this hypothesis, we examine the ECM of breast tumors by imaging Moffitt Cancer patients’ breast tumor samples using second harmonic generation (SHG) microscopy. The advantage of SHG imaging is that specific staining is not required and it can be done on IHC-stained slides to acquire expression data of proteins at the same time of fibrillar structures. We stained whole mount breast tumors samples with LAMP2b antibody (marker of acidosis and acid adapted cells (4, 39) and subsequently imaged those samples with bright field microscopy for LAMP2b and SHG for ECM and particularly collagen structure (Figure 2A). We found fibrillar structure inside the ducts and the location of the collagen correlates with the presence of acid adapted cells that also express LAMP2b.

**Figure 2:**
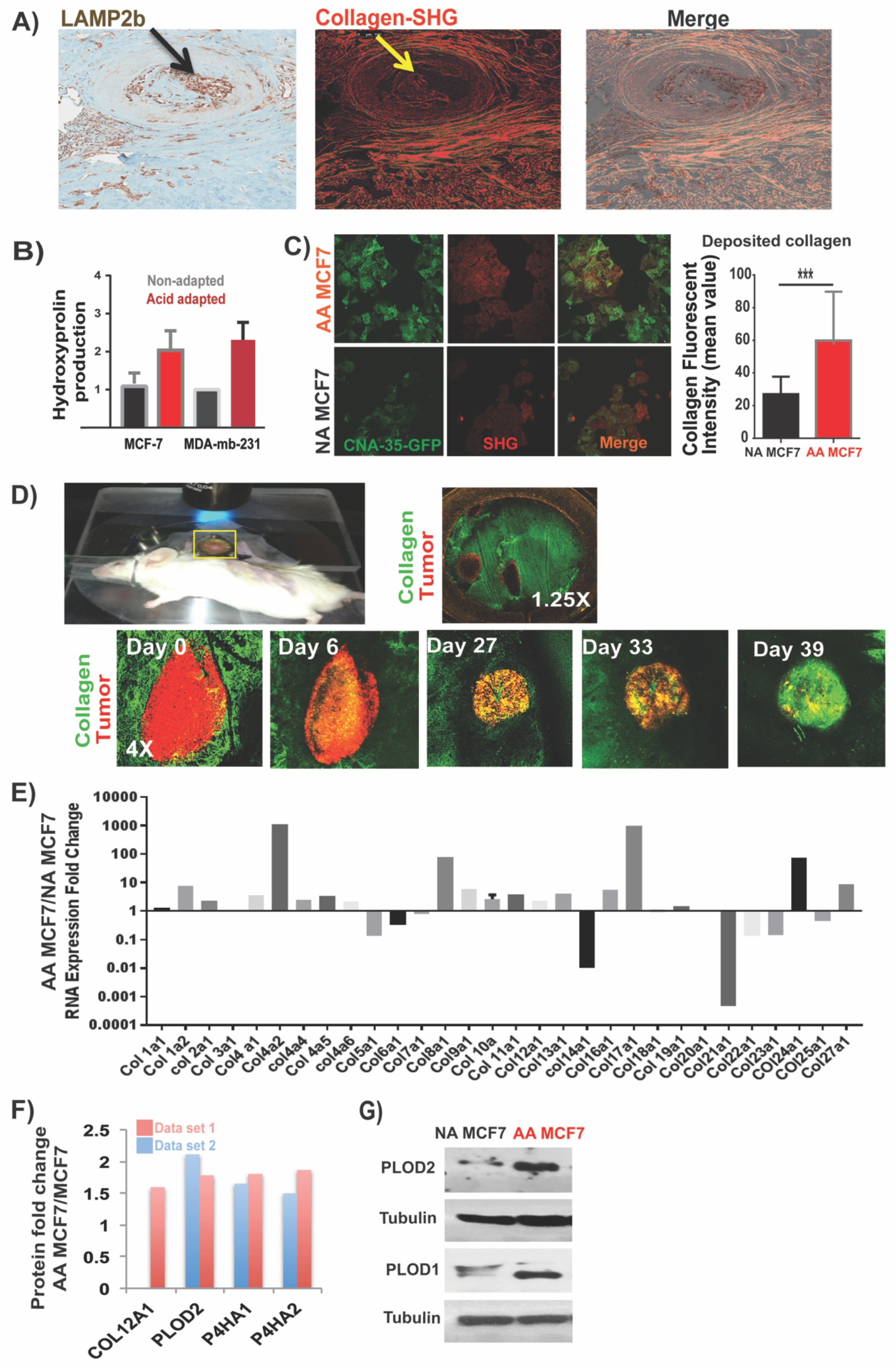
Acid adaptation promotes collagen production of cancer cells. A) Second Harmonic Generation (SHG) microscopy of breast tumors confirms the existence of collagen in the center of ducts in DCIS breast tumors. DCIS is the most acidic part of the tumor due to avascular nature of early carcinoma. B) Hydroxy proline assay showed higher amount of hydroxyprolin in acid adapted cancer cells compared to non-adapted ones. Hydroxyproline is an essential precursor of the collagen fibers that is necessary for its stability. The data is repeated in three biological replicates and data are presented as mean with SD as error bar. C) Fluorescent confocal microscopy coupled with second harmonic generation operator confirmed the higher expression of collagen and fibrillary structures in acid adapted cancer cells. Collagen production was measured using SHG and confocal microscopy on paraformaldehyde fixed breast cancer cells. CNA-35-GFP marker was used to distinguish collagen from other repetitive structure detected by SHG. Acid adapted cells produce more collagen significantly. Data is shown as standard deviation with mean as error bars with three separated biological replicates. D) Dorsal window chamber intravital microscopy of tumors in their matrix revealed collagen production of cancer cells. Tumor cells are marked with RFP and collagen is GFP. Clearly the green signal (Collagen-CNA-35-GFP) is increasing over time. E) Expression pattern of collagen genes in acid adapted MCF7 cells against non-adapted ones. The expression response pattern is quite heterogeneous and varies from over expression to lower expression. The y axis scale is in logarithmic some genes such as Col17a1 is highly expressed in acid adapted cells while Col21a1 is overly downregulated. F) SILAC proteomics analysis of AA MCF7 against NA MCF7 cells revealed increased expression of collagen production enzymes such as PLODs, and P4Has as well as some collagens in acid adapted MCF7 cells. G) Western blot validation of PLOD higher expression in acid adapted cells compared to non-adapted one. H) Secretome analysis of conditioned media of acid adapted cancer cells compared to non-adapted cells reveals higher amount of secreted TGM2 in both AA MCF-7 and AA MDA-mb-231. The experiment has three technical replicates and data is presented as mean with SD as error bar.

To further investigate collagen production by cancer cells and also the effect of acid adaptation on collagen production, we measured the hydroxyproline component of conditioned media from NA MCF7 and AA MCF7 as well as MDA-mb231 and AA MDA-mb-231 cancer cells. Hydroxyproline is a major amino acid in collagen that has a critical role in its stability. Hydroxyproline (4-hydroxyproline) is a non-proteinogenic amino acid that is the product of post-translational hydroxylation of proline. Because of high specificity of hydroxyproline to collagen, it can be used as an indicator of collagen content in isothermal conditions. Our result showed that the hydroxyproline content in conditioned media was increased in response of acid adaptation in both MCF7 and MDA-mb-231 cells. As cross validation of our findings *in vitro*, the collagen binding protein, CNA-35-GFP was then used to measure the collagen content of the cells and collagen deposited from AA MCF7 and NA MCF7 cells grown on glass bottom dishes. These same samples were also imaged with second harmonic imaging (SHG) which is also sensitive to regularized ECM structures (Figure 2C). This experiment additionally showed the higher expression of collagens in acid adapted cancer cells.

To study the collagen production and tumor acidosis, we also developed a dorsal window chamber (DWC) model to probe ECM and specifically collagen content of the tumors while growing (Figure 2D). DWC allows microscopic observation of the real time growth of tumors (40) and we have adapted this system to measure intra-and peri-tumoral pH (4, 41, 42). For DWC studies, we prepared spheroids of cancer cells, which were then implanted into the DWC and allowed to grow. The new live collagen imaging system was developed by applying CNA-35-GFP directly into the implanted tumor in the window chamber. Using this technique, we were able to track collagen structure changes in real time. We performed the experiment using two very aggressive acid producing and acid adapted cell lines (MDA-mb-231 and HCT-116) cells. Both of the cancer cells are marked with RFP protein and collagen binding protein is marked with GFP so we could image them both simultaneously. The pH of the whole window chamber area was measured by SNARF-1 (Seminaphtolrhodafluor-1, Life Technologies) as reported previously (4). Because of signal averaging and spatial kernelling, the ratiometric pH values were binned into four values, very low (< 6.7; black), moderately low (6.7-6.95; dark grey), neutral (6.85-7.1; light grey) and moderately high (>7.1; white) (Figure S1A and B). To study the relationship between pH and collagen structure, superposition of the acidity map from SNARF-1 with collagen images was used to correlate the structural differences of collagen extracted from the SHG images using Definiens software in different regions of the tumor with different pH_e_ values (Figure S1C). These analyses showed that collagen structure was not significantly different in more acidic areas of the tumor compared to non-acidic areas. Thus, an acidic environment does not appear to directly change the structure of collagens.

However, we observed accumulation of collagens inside the tumor area that rises over time parallel to increased acidification of tumor. Based on all of the above observations from different independent experiments we concluded that it is possible that cancer cells are producing collagen and the collagen production is promoted by acid adaptation.

To uncover what types of collagens are being induced by acid adaptation, we investigated the expression profile of 30 collagen genes in AA MCF7 and NA MCF7 using q-RT-PCR (Figure 2E). This showed a very heterogeneous response to acid adaptation in terms of collagen production. Some collagens such as Col17a1, Col4a2, Col8a1, Col9a1, Col10a1, Col11a1, and Col24a1 showed increased expression in acid adapted cells while Col5a1, Col6a1, Col7a1, Col14a1, and Col21a1 decreased. There were a group including Col3a1, Col18a1, and Col19a1 that showed no or very small change in their expression (Figure 2E). These results implied the different role of different collagens in response to chronic acidosis and their possibly differential role in the development and progression of tumors inside the duct.

To study different types of collagen overexpression in acid adapted cells at protein level, we searched our SILAC proteomics data on AA MCF7 versus MCF7 cells for matrix proteins and specifically collagen-related proteins. The proteomics data showed that collagen 12a1, and collagen modifying enzymes, such as PLODs and P4HAs, were increased in acid adapted cells (Figure 2F). Notably, Col12a1 mRNA was not significantly upregulated, suggesting translational-level control. PLOD proteins are membrane-bound homodimeric enzymes localized to the cisternae of the rough endoplasmic reticulum and catalyze the hydroxylation of lysyl residues in collagen peptides. These hydroxylysyl groups are different carbohydrates attachment sites that are critical for the stability of collagen intermolecular crosslinks. P4HA proteins are components of prolyl 4-hydroxylase, a key enzyme in the thermostabilization of collagen. Prolyl 4-hydroxylase catalyzes the formation of 4-hydroxyproline, which is necessary to the proper three-dimensional folding of newly synthesized procollagen peptides. We validated the higher production of PLOD1 and PLOD2 in acid adapted cells with western blotting (Figure 2G). We conclude collagen production is elevated in acid adapted cancer cells.

### Acid adapted cells use collagen remodeling enzymes to engineer the niche

In our proteomics data (4) we also observed that Transglutaminase 2 (TGM2) had the second highest fold change in acid adapted compared to non-adapted MCF7 cells (Figure 3A). TGM2 is a calcium-dependent enzyme of the protein-glutamine γ-glutamyltransferases family that makes inter-and/or intramolecular bonds that renders proteins resistant to proteolysis degradation. TGM2 is found both in the intracellular and the extracellular spaces of cells and plays different roles in different spaces. The extracellular form binds to proteins in ECM such as collagen and fibronectin and promotes cell adhesion, ECM stabilization, wound healing, and cellular motility (43–45). Thus, TGM2 may enable the acid adapted cancer cells to engineer and change the ECM by even more advanced mechanisms than collagen deposition.

**Figure 3.**
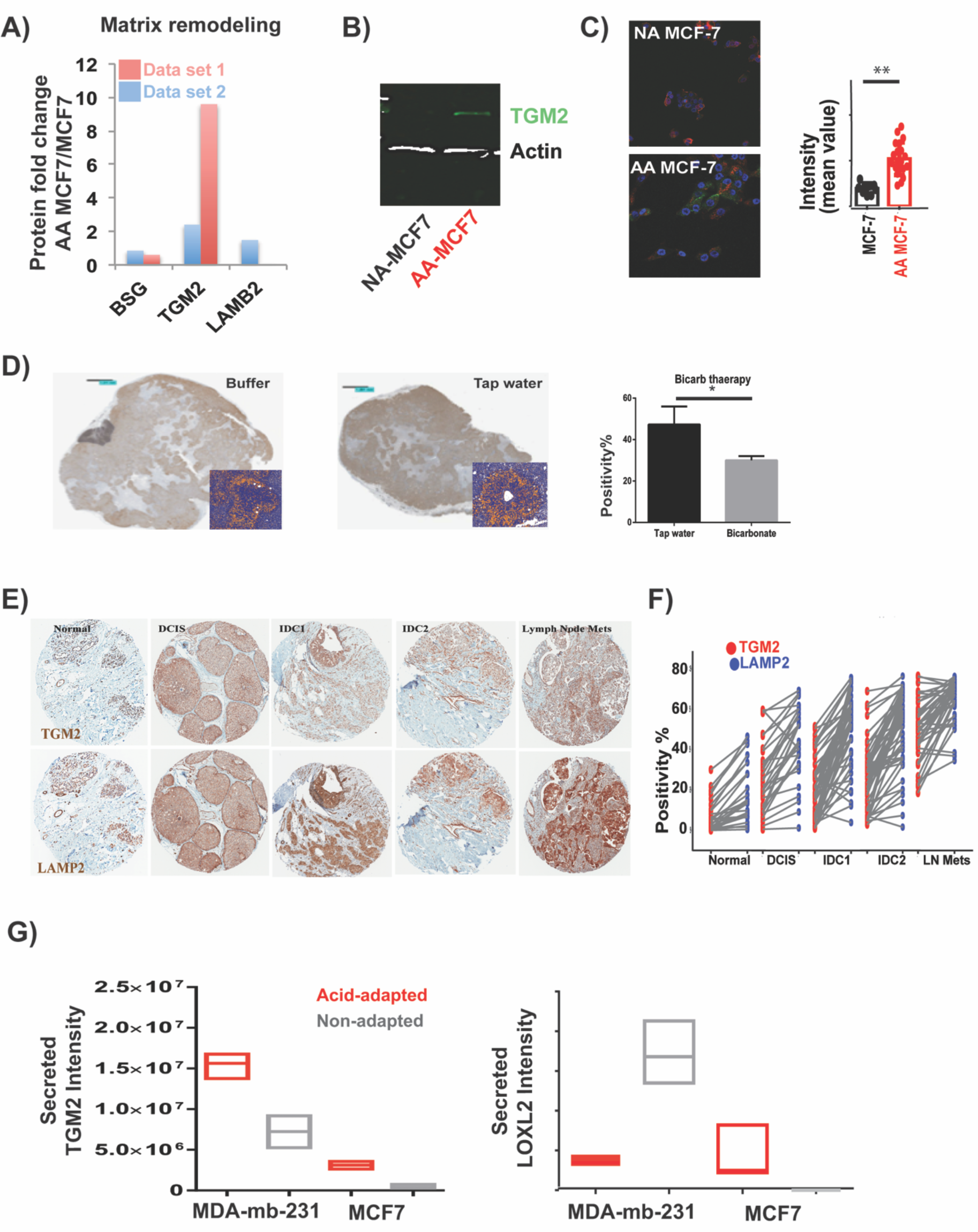
Acid adapted cells use collagen remodeling enzymes to engineer their niche. **A)** TGM2 validation by **B)** western blot and **C)** Immunocytochemistry (ICC). TGM2 expression in acid-adapted cancer cells is significantly higher compared to non-adapted cells. **D)** Bicarbonate buffer therapy in animal reduced the expression of TGM2 compared to control group, indicating the effect of acidosis on expression of TGM2. **E, F)** Translation of TGM2 expression to Clinique and patient samples. LAMP2b is a marker of acidosis that we reported in our previous work. The TMA from same patients was stained for TGM2 and expression level of TGM2 and LAMP2b was compared for each sample. There is a correlation between LAMP2b and TGM2 expression in patients. **G)** Acid adapted MCF-7 cells not only have higher amount of TGM2 and LOXL2, but also secret more of these enzymes to the environment. Both enzymes have been shown to play role in collagen crosslinking and stability. This strengthen our acid-induced niche engineering phenotype of cancer cells that they build the niche they need it to survive the harsh environment such as acidosis.

To validate the overexpression of TGM2 in our acid adapted cells, we performed western blots and Immunocytochemistry (ICC) on acid-adapted and non-adapted MCF-7 cancer cells. (Figure 3B, and 3C). Then, we showed in mice that we can reduce the expression of TGM2 using bicarbonate buffer consistent with our hypothesis about the acid-induced nature of this protein (Figure 3D). To investigate this in patient samples, we compared the expression of TGM2 and LAMP2b, previously proven as a marker of acidosis in solid tumors in human cancers, in each patient. The expression of TGM2 and LAMP2b was correlated in 240 different patients’ samples of different stages from adjacent normal to DICS, IDC 1, IDC 2, and Macrometastasis (Figure 3E and 3F). We found that TGM2 expression is elevated by acid in tumors; however, as we discussed above, the localization of this enzyme is highly related to its activity. The protein crosslinking and niche engineering activity is assigned to extracellular TGM2. To determine if TGM2 is actually secreted to the extracellular milieu and its extracellular expression is increased in acid-adapted cells we did secretome analysis of MCF-7 and AA-MCF7 using mass spectrometry-based proteomics. The secretome analysis showed the higher expression of TGM2 in both AA MCF-7 and AA MDA-mb-231compared to the corresponding non-adapted cells (Figure 3G). We also observed other crosslinking enzymes such as LOXL2, an enzyme that has been shown to play a role in collagen crosslinking and stability (46), are over expressed in AA MCF7 (Figure 3G). Western blot analysis validated overexpression of LOXL2 in AAMCF7 cells (Figure S4) similar to TGM2. The MDA-mb-231 cells look as if they only use TGM2 secretion to crosslink the collagen.

To further test our hypothesis in cancer patient samples we stained for TGM2 in whole mount breast tumors. The results showed higher amount of TGM2 in acidic regions such as center of DCIS and also co-registered with collagen and other fibers in ECM and extracellular spaces (Figure S6). We also looked at the expression level of Matrix MetaloProteinases (MMPs) in our secretome data and found no overexpression of these proteins in the extracellular spaces of the cells (Figure S7A). To validate these results and also validate the activity of MMPs in acid-adapted and non-adapted cells we also conducted CQ-collagen assay and found no difference (Figure S7B).

The above findings imply that adaptation to an acidic environment by cancer cells induces matrix remodeling, which can be viewed as a type of “niche engineering” to promote survival.

### Acid-induced collagen remodeling is controlled by SMAD proteins and Ras activity

Collagen production is commonly under the control of TGF-β signaling through SMAD proteins. Further, Ras signaling plays a prominent role in the conversion of TGF-β signaling from anti-to pro-oncogenic and thus promotes tumor progression (47). However, the mechanisms of the synergy between oncogenic RAS and TGF-β signaling are enigmatic. It has been shown that Ras activation can switch TGF-β family function from tumor-suppressive to tumor-promoting functions that will increase tumor growth and early dissemination of cancer cells (48). It also has been shown that in cardiac myocytes extracellular acidosis can activate the Ras and ERK1/2 (49). We hypothesized that this pathway can be hijacked by cancer cells to activate Ras through extracellular acidosis, and that this will induce SMAD proteins to produce essential collagens for tumor cell survival. To test this hypothesis, we investigated the nuclear expression of SMAD4 in acid adapted and non-adapted MCF7 cells, as nuclear localization is a marker for SMAD activity. The nuclear expression of SMAD4 was higher in AA MCF7 cells compared to NA MCF7 while the cytoplasmic expression of SMAD4 in NA MCF7 was higher (Figure 4A and 4B). To study the possible role of SMAD3 in translocation of SMAD4, we investigated nuclear and cytoplasmic expression of pSMAD3 in AA MCF7 and MCF7 cells using ICC and confocal microscopy. The nuclear expression of pSMAD3 was significantly higher in AA MCF7 cells and cytoplasmic expression was lower (Figure 4C and 4D). As a control we also looked at SMAD2 expression in both cells and found no differences in nuclear or cytoplasmic expression. Then we investigated the activity of Ras pathway in acid adapted MCF7 and MCF10AT cells compared to their non-adapted counterparts. We observed that expression of K-Ras was higher in acid adapted cells while N-Ras and H-Ras were downregulated. To investigate if Ras is actually active in acid adapted cells, we measured the active Ras isoform using active Ras pull down. Active Ras was significantly higher in both acid-adapted MCF7 and MCF10AT compared to their non-adapted counterparts (Figure 4E). When Ras is active it will activate ERK through a signaling cascade driven by phosphorylation. pERK will phosphorylate SMAD3 through SMAD1 that eventually translocate SMAD4 to the nucleus and activate collagen production genes. To support this sequence of events, we performed western blot and observed that ERK1/2 is phosphorylated and active in acid-adapted cells (Figure 4E). To validate the effect of acid-induced activation of Ras in vivo we injected the acid adapted (activated Ras) and non-adapted MCF10AT cells into flanks of SCID mice. Acid-adapted MCF10AT tumors grew much faster and to a larger size compared to non-adapted ones (Figure 4F). Based on our findings, we propose a role for acid-adaptation activation of the Ras pathway to mobilize SMADs for collagen production (Figure 4F).

**Figure 4.**
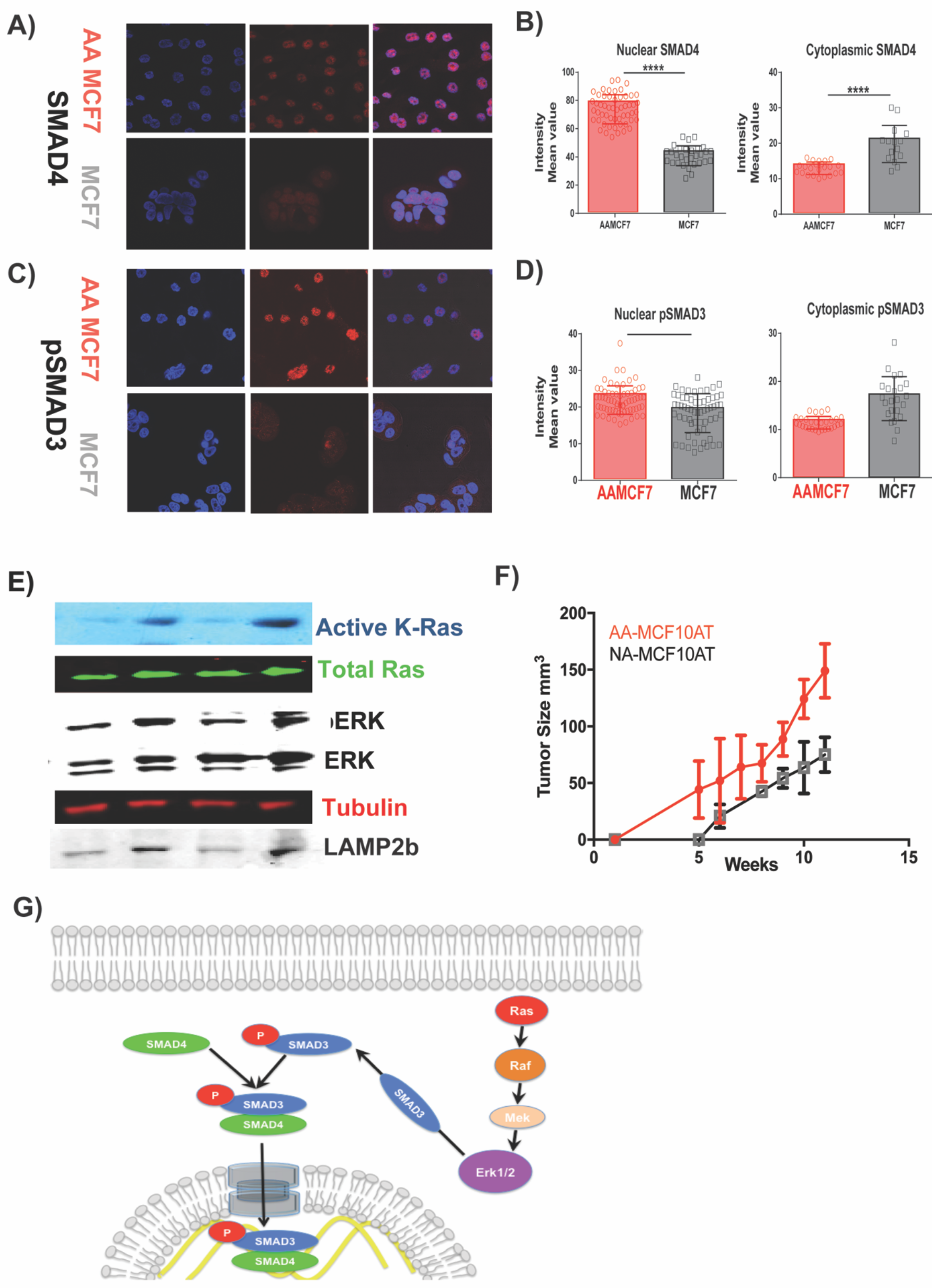
Acid-induced collagen production is controlled by SMADs and K-Ras. A) Immunocytochemistry (ICC) of acid adapted MCF7 and non-adapted MCF7 for SMAD4. SMAD4 is a transcription factor that controls the production of collagen when it is located in nucleus. B) Nuclear localization of SMAD4 versus its cytoplasmic localization revealed the higher nuclear localization of this protein in acid adapted cells. C) pSMAD3 staining of acid-adapted and non-adapted MCF7 cells showed upregulation of pSMAD3 in acid-adapted cells. D) Nuclear and cytoplasmic localization analysis of pSMAD3. SMAD3 is a cytoplasmic protein that binds to SMAD4 in its phosphorylated form and translocates SMAD4 into the nucleus. pSMAD3 has higher nuclear expression and lower cytoplasmic expression in acid-adapted MCF7 cells compared to non-adapted MCF7. E) Ras activation analysis and downstream protein regulations. K-Ras is activated in acid-adapted cancer cells that activate ERK through phosphorylation. F) Tumor growth of acid adapted MCF10AT and non-adapted ones injected to nude mice. AA MCF10AT cells frequency of forming tumor is higher and the tumors that are formed grow faster. G) Schematic of Ras activation role in collagen production in cancer cells. We proposed Ras activation phosphorylate ERK that can activate SMAD3 through phosphorylation. pSMAD3 will translocate SMAD4 to nucleus to activate collagen production genes.

### The role of collagens in cancer cell fitness and evolution

As part of our cross-validation analysis of these findings in an independent dataset, we investigated the expression of collagens in breast tumors compared to their adjacent normal cells in the TCGA. Collagens are heterogeneously expressed in tumors and have different effects on overall patients’ survival (Figure 5A). Many of the collagens that are over-expressed in tumors compared to normal cells are rare collagens such as Col10a1 and Col11a1. It struck us that cancer cells may use these rare collagens because it provides them some advantages over normal cells that normally don’t express these proteins. Further we wondered if this overexpression is also seen in our acid adapted cells. Therefore, we compared the genes from TCGA data (Figure 5A) with our genes that have higher expression in cells grown in acid media (Figure 5B). Then, we made a table of the genes that are either high or low in both TCGA data and acid adapted cells (Figure 5C). This analysis can give us information about the genes playing a role both in acid adaptation and tumor evolution. We hypothesize that, if these genes play a role in tumor progression and evolution and more precisely acid-induced tumor progression, then any mutation in those genes must affect the cancer cells’ fitness and could have a positive effect on patients’ survival. To test this hypothesis, we categorized the genes from figure 5C in two groups. First, the genes expressed at high levels both in acid pH and tumors from TCGA: (Col1a2, Col2a1, Col4a5, Col8a1, Col9a1, Col10a1, Col11a1, Col13a1, and Col24a1), which we suspect are tumor promoting “bad” collagens. Second, the collagens that are lower in both cells grown in acidic media and TCGA tumors: (Col4a6, Col6a1, Col7a1, Col14a1, Col15a1, Col16a1, Col21a1, Col23a1, and Col25a1), which we suspect have anti-tumor effects, or are “good” collagens. Thus, mutations in “bad” collagens should promote patient survival and vice-versa. Figure 5D shows that, if patients have mutations in “bad” collagen genes, their survival is higher and vice versa, if a patient has mutation in “good” collagens, their survival was lower. Thus, the overall model states that cancer cells under acidic conditions that can express these gene products can adapt better to stress, acidosis in this case, by constructing and engineering a niche to promote their survival. Any mutations that reduce this ability will affect the cancer cells’ fitness and viability that eventually will affect the patients’ survival. This is a good area for mechanistic follow up studies and a direction for our future works.

**Figure 5.**
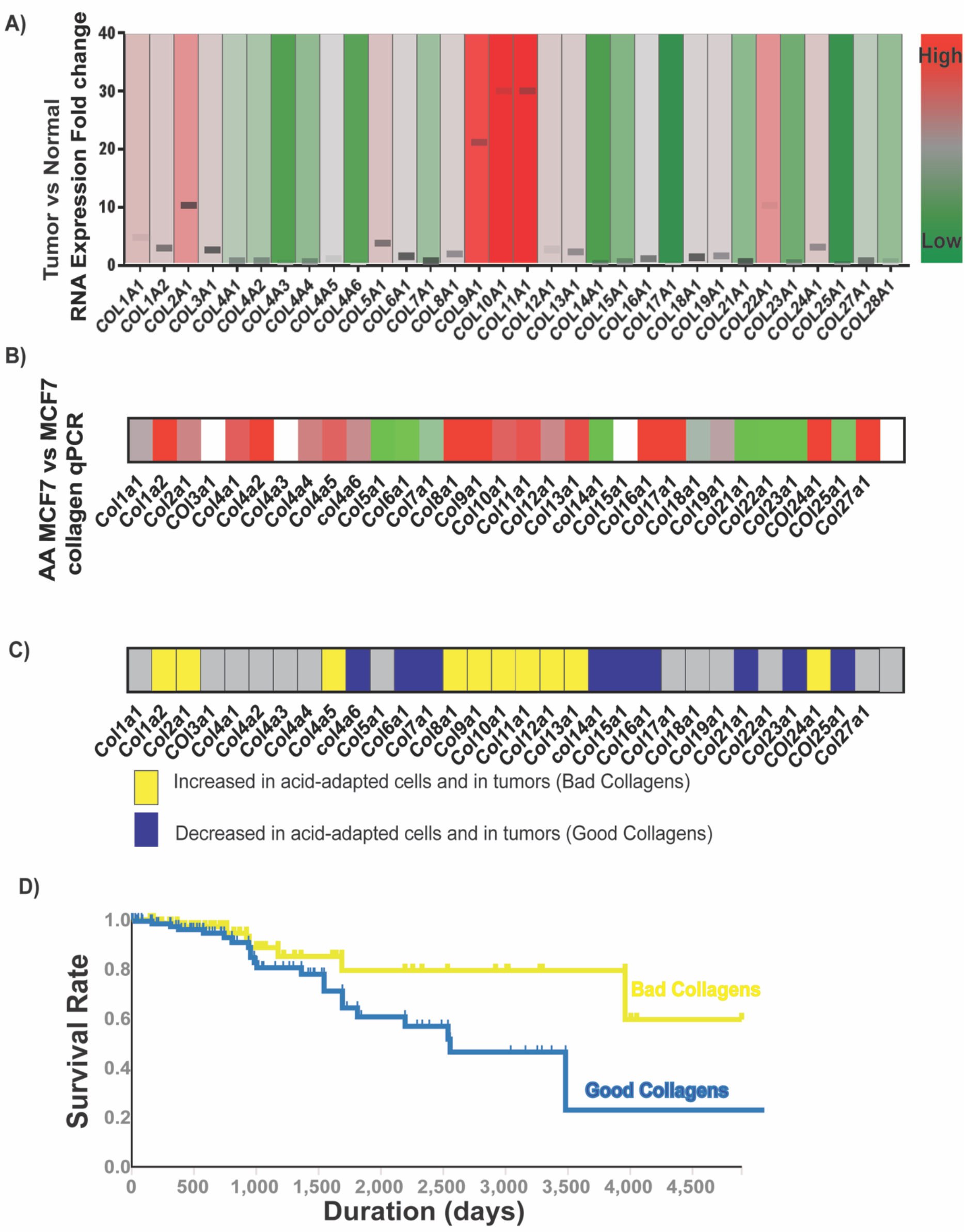
Collagen production of cancer cells in patient samples. A) TCGA data analysis of different collagen expression in tumor cells compared to their adjacent normal. There is a heterogeneity in collagens expression. The columns are color coded as red highest expression and green the lowest. B) The same color coding as A for the q-PCR data of acid adapted versus non-adapted MCF7 cells. The ones that have higher expression are correlated with acid-induced ones such as Col10a1 and Col11a1. C) Comparison of TCGA data from A with B, the over expressed collagens in acid adapted cells. The yellow is the gene that is over expressed both in A and B, Blue is low in both A and B, and grey is neither. This correlation can be used to find the collagen genes that play role in acid-induced evolution of cancer cells D) Survival analysis of patients with mutation in different collagen genes. Based on the expression level analysis showed in C, collagens were categorized in two groups: High expressing collagens in both patients and acid pH (bad collagens) and low expressing collagens (good collagens). The overall survival of patient with bad collagen mutations is higher than patient with mutations in good collagen genes.

## Discussion

### Collagen production and engineering is necessary for early stage cancer cells survival

Cancer is a dynamic system composed of cancer cells and their microenvironment that is always changing. Interactions among components of the cancer ecosystem shapes the forces driving tumor evolution. Evolution of cancer cells in their local microenvironment is governed by their niche contents that creates evolutionary selection forces. Tumor cells are able to alter the local environment or even build a new environment to promote their viability and growth in any emerging new environment. This phenomenon is known as "niche construction and engineering" and we believe it to be a key part of the cancer cells self-defined fitness strategies. Each subtle change in tumor niche, from cancer or normal cells or stroma cells, can have profound effects on the tumor host interaction that will affect the tumor growth dynamics dramatically. We have recently shown that different sub-populations in tumors engineer the local ecology to favor its own growth and survival (35).

The metabolic and niche remodeling changes in early cancers are components of a non-linear evolutionary dynamics, wherein the changing microenvironment selects for phenotypes that, in turn, alter the microenvironment in ways that provide subsequent selection forces for subsequent phenotypes. Like organisms, cancer cells frequently modify local resource distributions, influencing both their microenvironments and the evolution of traits to improve fitness (50). In ecology, these processes are known as “niche engineering”, wherein one species will modify its habitat to maximize its own fitness, often to the detriment of other species within the same habitat. Examples in nature include ants and termites who construct their habitats that are the new source of selection of their own species (51–53). The best example of niche engineering in cancer cells, and the most well-studied one to date, is the pre-metastatic niche in lymph nodes, which is thought to be mediated by tumor-derived exosomes (54). Niche construction strategies of early cancer are relatively understudied.

In this paper we show the reciprocal interaction of tumor cells and their microenvironment focusing on adaptive strategies of cancer cells in their harsh acidic environment. We found niche construction and engineering as an evolutionary adaptive strategy exploited by cancer cells to survive in emerging harsh environment that also lead them to evolve and develop into new stages. We see the adaptation of cancer cell to acidosis as a self-defined fitness function. Acid adapted cell survival and proliferation are determined entirely by their own heritable phenotypic properties i.e. cells can develop independence from normal tissue control through either mutations or phenotypic plasticity and reversible adaptation that disrupt their response to the host signals. A self-defined fitness function through plasticity or adaptation is similar to when tissue control signals are lost due to inflammation or infection (55). Here we also show how adaptation to microenvironmental selection pressure such as acidosis regulates cancer cells self-defined fitness as part of their evolutionary strategies to survive and invade, in this case to construct and engineer their favorable niche for growth and proliferation. Here we showed that over expressing some collagen genes and down regulation of the others can give the cancer cells the self-defined fitness function that will help them reconstruct and engineer their niche in their own favor to survive, proliferate or even metastasize later.

In this paper we showed for the first time the molecular biology mechanism behind the acid-induced niche reconstruction and engineering that was also related to an evolutionary subject of self-defined fitness. We believe evolution is theory of cancer and understanding all the evolutionary disciplines ruling the cancer growth and development will help us fighting against cancer and come up with strategies to control it. Figure 6 shows a schematic of breast cancer tumors (as representative of many epithelial solid tumors) growth regarding the acidic microenvironment and collagen morphology and structure change. Solid tumors are profoundly and continuously acidic due to Pasteur and Warburg effect. The collagen morphology variation has been reported previously with mostly unknown trigger and mechanism. Here we show how microenvironmental factors such as acidosis can promote the collagen morphology changes over cancer progress. Acid adapted cells gain the ability of producing the extra cellular matrix component such as collagen that help them to survive and progress. All the above described changes are summarized in a phenotype: niche construction and engineering ability, and as we showed in this paper any mutation changing this phenotype can hurt the cancer cells and benefit patients.

**Figure 6.**
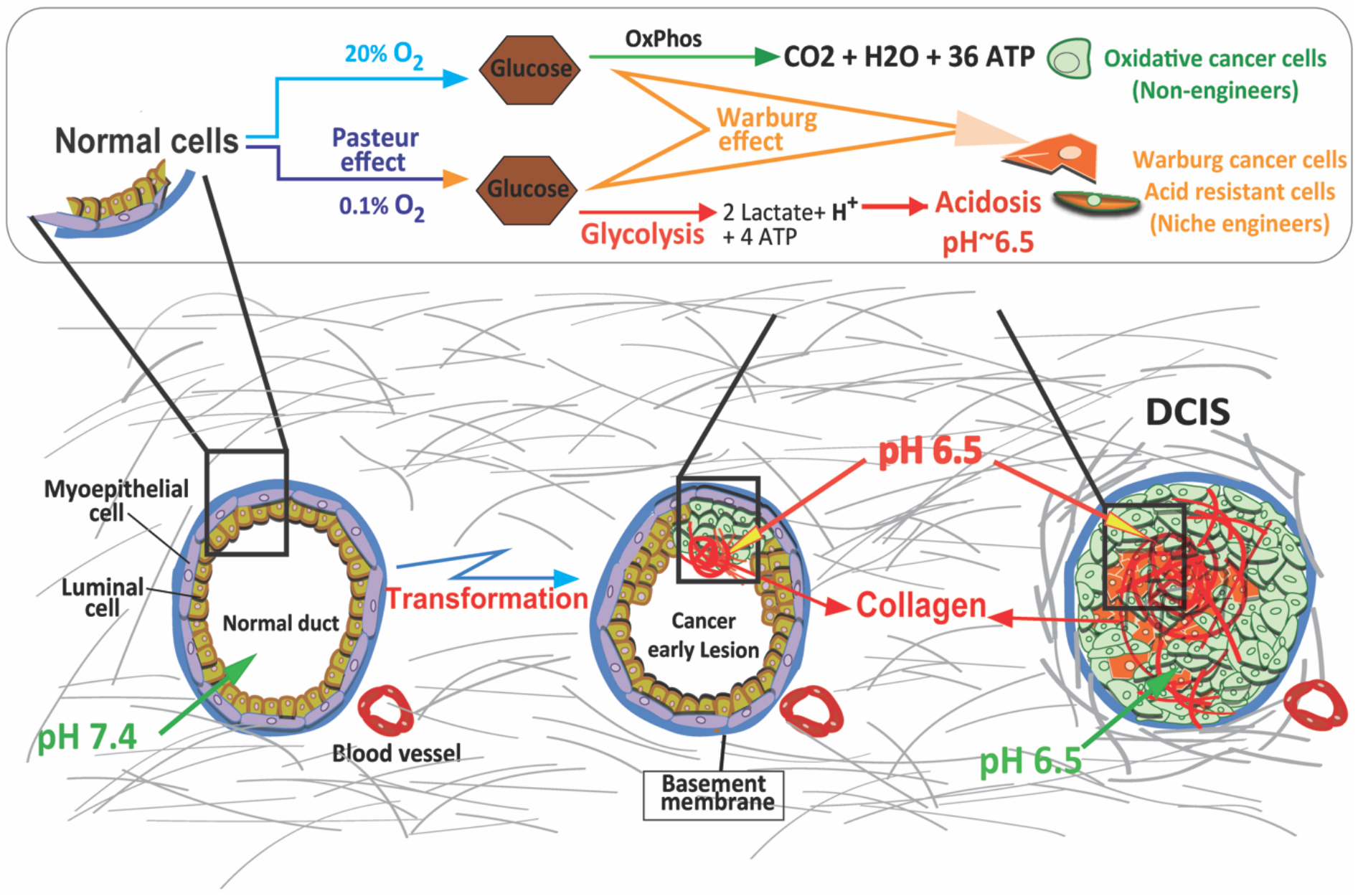
Acid induced collagen production promote tumors early stage development and progress in different cancer models. Schematic of breast cancer tumors growth regarding the acidic microenvironment and collagen morphology and structure change. Solid tumors are profoundly and continuously acidic due to Pasteur and Warburg effect. The collagen morphology variation has been reported previously with mostly unknown trigger and mechanism. Here we show how microenvironmental factors such as acidosis can promote the collagen production and its role in tumor evolution. At the early stages, acid-adapted cancer cells inside the DCIS gain the ability of producing the extra cellular matrix component such as collagen. This will enable them and probably their neighbors surviving the anoikies and progress toward next stage.

## Acknowledgments

We thank the staff of the Molecular Genomics Core, Proteomics and Metabolomics Core, and Analytical Microscopy Core of the Moffitt Cancer Center for their assistance with these studies. The Cores are partially funded by the National Cancer Institute through P30CA076292 as a Cancer Center Support Grant. M.D. and R.J.G. was supported by grants from NCI (R01CA077571) and NIH/NCI PSOC(1U54CA193489).

## Author Contributions

M.D. conceived the study, developed the methodology, designed and performed experiments and computational analysis. L.X., N.T., and T.C.performed the experimental work. J.J. supervised the microscopy analysis work. M.D., B.F. and J.M.K. performed the proteomics and its data analysis. A.K. and M.D. performed TCGA data analysis. N. D.G. supervised the spinning disk experimental work. All authors participated in the interpretation of the results. M.D. wrote the paper with contributions from all authors. All authors have read and approved the final version of the manuscript. M.D. directed the study.

## Method

### Cell culture and *in-vitro* acid adaptation

MCF-7, MCF10-AT, MDA-mb-231, and MCF10A cells were acquired from American Type Culture Collection (ATCC, Manassas, VA, 2007–2010) and were maintained in DMEM-F12 (Life Technologies) supplemented with 10% fetal bovine serum (HyClone Laboratories). Growth medium was further supplemented with 25□mmol□l^−1^ each of PIPES and HEPES and the pH adjusted to 7.4 or 6.7. Cells were tested for mycoplasma contamination and authenticated using short tandem repeat DNA typing according to ATCC’s guidelines. For acute acidosis cells were exposed to acidic media for 72 hours. To achieve acid adaptation, cells were chronically cultured and passaged directly in pH 6.7 medium for more than 3 months.

### SILAC labelling and Proteomics

This technique was described in details in our previous publication (4).

### Secretome proteomics

Acid adapted and non adapted cancer cells were cultured in 10 cm dishes at 50% confluency. After 24 hours, their media was replaced with serum free media, which was collected after 12 hours of incubation and lyophilized. Proteins were dissolved in denaturing lysis buffer containing 8M urea, 20 mM HEPES (pH 8), 1 mM sodium orthovanadate, 2.5 mM sodium pyrophosphate, and 1 mM β-glycerophosphate. A Bradford assay was carried out to determine the protein concentration. The proteins were reduced with 4.5 mM DTT and alkylated with 10 mM iodoacetamide. Trypsin digestion was carried out at room temperature overnight, and tryptic peptides were then acidified with 1% trifluoroacetic acid (TFA) and desalted with C18 Sep-Pak cartridges. LC-MS/MS analysis was carried out with a nanoflow ultra high performance liquid chromatograph (RSLC, Dionex, Sunnyvale, CA) coupled to an electrospray mass spectrometer (Q-Exactive Plus, Thermo, San Jose, CA). The sample was first loaded onto a pre-column (2 cm × 100 µm ID packed with C18 PepMap100 reversed-phase resin, 5µm particle size, 100Å pore size) and washed for 8 minutes with aqueous 2% acetonitrile and 0.04% trifluoroacetic acid. The trapped peptides were eluted onto the analytical column, (C18 PepMap100, 75 µm ID × 25 cm, 2 µm particle size, 100Å pore size, Dionex, Sunnyvale, CA). The gradient was programmed as: 95% solvent A (2% acetonitrile + 0.1% formic acid) for 8 minutes, solvent B (90% acetonitrile + 0.1% formic acid) from 5% to 38.5% in 60 minutes, then solvent B from 50% to 90% B in 7 minutes and held at 90% for 5 minutes, followed by solvent B from 90% to 5% in 1 minute and re-equilibration for 10 minutes. The flow rate on analytical column was 300 nl/min. Sixteen tandem mass spectra were collected in a data-dependent manner following each survey scan using 15 second exclusion for previously sampled peptide peaks. MS1 resolution was set at 70,000 and MS/MS resolution was set at 17,500 with ion accumulation (maxIT) set to 50 ms. MaxQuant (version 1.2.2.5) was used to identify and quantify the proteins.

### Western blotting

Acid-adapted and non-adapted MCF10-AT, MCF-7, and MDA-MB-231 cells were grown with the same number of passages and used for whole-protein extraction. Lysates were collected RIPA buffer containing 1 × protease inhibitor cocktail (P8340; Sigma-Aldrich). Twenty micrograms of protein per sample was loaded on polyacrylamide–SDS gels, which later were electrophoretically transferred to nitrocellulose. Membranes were incubated with primary antibodies against rabbit polyclonal LAMP2 (1:1,000, ab18529 Abcam), mouse mono clonal PLOD2 (1:1,000, Fischer Scientific), Rabbit poly clonal PLOD1(<u>LS-C163796-400</u>), K-Ras, N-Ras, and H-Ras (Santa Cruz Biotechnology) and GAPDH (1:4,000, antirabbit; Santa Cruz Biotechnology).

### Immunofluorescence

Cells cultured at pH 6.7 chronically and pH 7.4 of with the same number of passages were rinsed with PBS, fixed in cold 4% paraformaldehyde for half an hour and then blocked with 4% bovine serum albumin in PBS. Samples were incubated with LAMP2 rabbit polyclonal primary antibody (1:100; ab 37024 Abcam) and secondary Alexa-Fluor 488 antirabbit (1:500) antibody. Coverslips were mounted using ProLong Gold Antifade Reagent (Life Technologies) and images were captured with a Leica TCS SP5 (Leica) confocal microscope.

### Spinning disk cell adhesion measurement

To measure the strength of cellular adhesion, a hydrodynamic flow chamber is filled with a fluid of known viscosity and density (56, 57). Inserted into the chamber is a spinning shaft that holds via vacuum a glass cover slip on which cells have adhered. The cells are therefore exposed to the shear force of the laminar flow at a given rotational speed. The shear stress varies linearly with radius:

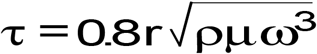

where *r* is the radial position along the substrate, *μ* is the viscosity, *ρ* is the density of the solution, and *ω* is the angular velocity. After spinning for 5 min, the remaining adherent cells were fixed in 3.7% formaldehyde, permeabilized with 0.1% Triton X-100, and stained with Hoechst dye to identify the nucleus. The number of adherent cells was counted at specific radial positions using an Eclipse Ti-U fluorescent microscope (Nikon Instruments, Melville, NY) fitted with a motorized stage and NIS-Elements Advanced Research software (Nikon Instruments). Sixty-one fields were analyzed per substrate and the number of cells at specific radial locations was then normalized to the number of cells at the center of the substrate where negligible shear stress was applied to calculate the fraction of adherent cells *f*. The detachment profile (*f* versus *τ*) was then fit with a sigmoid curve

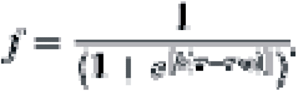

The shear stress for 50% detachment (τ_50_) was used as the mean cell-adhesion strength.

### Spheroid assay

Perfecta3®96-Well Hanging Drop Plates and low attachment U shape plates were used to grow the primary spheres containing 10000 cells (acid adapted MCF-7 or non-adapted MCF-7) without using any matrix such as Matrigel or collagen. After 24 hours the spheres media were changed to acid or normal media. Acid-adapted MCF7 cells grow robust spheres around day7 while NA MCF7 never grow real sphere. Incucyte microscope was used to image the sphere growth. Data were analyzed using custom built Incucyte software.

### Soft agar assay

Soft–agar colony formation assay is used to study the anchorage–independent growth of tumor cells. 1 million NA and AA MCF7 cells are trypsinized, counted and dissolved in 1 ml DMEM with 10% FBS. 4% agar was melted by microwave and kept in 56 C water bath to be used as bottom layer. To prepare the bottom layer 4:1 ratio of 4% agar: DMEM media with 10% serum was mixed and poured in 6 well plates (1 ml per well). The plates were placed in hood to let the gel solidify. For top layer 0.4% agar gel was made by adding the DMEM medium with 10% serum. 100 cells were added per milliliter of the mixture and added on top of the bottom layer. Plates were placed in 37C incubator for 2-3 weeks and colonies were counted by microscopy or crystal violet staining.

### Dorsal window chamber

Intravital multiphoton confocal microscopy was used to create a pH map using SNARF-1, a ratiometric pH-sensitive fluorescent probe that has altered emission wavelengths at basic and acidic pH values, and to observe collagen dynamics over time *in vivo*. Tumour constructs were prepared using the tumour droplet method as described previously (4, 41). HCT-116-RFP or MDA-MB-231-RFP cells were suspended in 0.8□mg□ml^−1^ of type-1 collagen (BD Bioscience #354249) and 1 × DMEM at a final concentration of 2.5 × 10^6^ cells□ml^−1^. Then, a 48-well non-tissue culture plate was used to place 5–15-μl drops (depends on the size of the tumour) of the tumour suspension in it. The droplets were polymerized in the center of the well after 20–30□min of incubation at 37□°C. Next, 25□μl of 1.25□mg□ml^−1^ collagen 1 was added to the surrounding of the tumour droplets. This helped the tumour droplets to have nicely defined borders. Then 200□μl of growth medium (DMEM/F12 with 10% fetal bovine serum) was added to the wells and incubated in 37□°C.

Meanwhile, a DWC was prepared to implant into the recipient mice for next 24–48□h of culturing the tumour droplet constructs. Constructs were added aseptically into the wound area right after the surgery. Intravital microscopy was started from 24–48□h after surgery to acquire tumour integrity and growth. To develop a pH map for each tumour and its microenvironment inside the chamber, we tail-vain-injected SNARF (a fluorescent molecule that has different emission in basic and acidic pHs) to the mice and imaged after 30□min. For SNARF imaging, we used excitation with an Argon laser tuned to 543□nm and emission was collected in 590–640□nm bandpass filter. Images were collected using an Olympus FV1000 MPE (multiphoton) microscope. Analysis was carried on using Image-Pro Plus v6.2 (Media Cybernetics; Bethesda, MD).

To be able to observe collagen structure in window area we developed our method by adding CNA-35-GFP (100 uM) directly to the window area for 10 minutes followed by 3 times washes with physiological saline to remove the excess marker. The marker was renewed every week by removing the window coverslip and repeating the above process. The collagen images were acquired by 488 nm excitation bandwidth.

### Active Ras measurement

Ras superfamily of small GTPases are active when bound to GTP and inactive when the triphosphate is hydrolyzed to GDP. The Active GTPase pull-down and Detection Kits enriches Ras active form using a GST-protein binding-domain fusion that is selective for active Ras. The amount of enriched active Ras is measured using specific antibody and western blotting. We used Active Ras Detection Kit #8821 from cell signaling technology that uses GST-Raf1-RBD fusion protein to bind the activated form of GTP-bound Ras, which can then be immunoprecipitated with glutathione resin. Ras activation levels are then determined in western using a Ras mouse mAb.

### Animal experiment

All animals were maintained in accordance with IACUC standards of care in pathogen-free rooms, in the Moffitt Cancer Center and Research Institute (Tampa, FL) Vivarium. One week before inoculation with tumour cells in the mammary fat pads, female nu/nu mice 6–8 weeks old (Charles River Laboratories) were placed in two cohorts per experiment randomly selected. Acid adapted and non-adapted MCF7 and MCF10AT cells were mixed with Matrigel in cold PBS and was injected into mammary fat pad of animals. The tumor size was monitored by caliper three times a week and by Ultrasound every week. When the tumor reached 1500 mm^3^ of size, animals were humanely killed and tumours were extracted, fixed in 10% formalin, paraffin embedded and further processed for IHC.

### Microarray analysis

Affymetrix expression data for ColXa1 and Col11a1 genes in patient samples were produced from publicly available data sets of Moffitt Cancer Center patients. The CEL files for the tumor samples were downloaded from the Gene Expression Omnibus (GEO) database (http://www.ncbi.nlm.nih.gov/geo/), data series GSE2109. Normal tissue data were from the GEO data series GSE7307, Human Body Index. The CEL files were processed and analyzed using the MAS 5.0 algorithm (Affymetrix) and screened through a rigorous quality control panel to remove samples with a low percentage of probe sets called present by the MAS 5 algorithm, indicating problems with the amplification process or poor sample quality; high scaling factors, indicating poor transcript abundance during hybridization; and poor 30/50 ratios, indicating RNA degradation either before or during processing. The remaining samples were normalized to the trimmed average of 500 in the MAS 5 algorithm before comparison of the expression values across tumours and normal samples.

### Immunohistochemistry

For human tissues, a TMA containing formalin-fixed and paraffin-embedded human breast tissue specimens was constructed in Moffitt Cancer Center histology core. The TMA contains 27 normal breast tissue, 30 DCIS, 48 invasive ductal carcinomas without metastasis, 49 invasive ductal carcinomas with metastasis and 48 lymph node macrometastases of breast cancer. Cores were selected from viable tumour regions and did not contain necrosis. A 1:400 dilution of anti-LAMP2b (#ab18529, Abcam), was used as primary antibody. Normal placenta was used as a positive control for LAMP2 and for the negative control, an adjacent section of the same tissue was stained without application of primary antibody, and any stain pattern observed was considered as nonspecific binding of the secondary. Immunohistochemical analysis was conducted using digitally scanning slides. The scoring method used by the pathologist reviewer to determine (1) the degree of positivity scored the positivity of each sample ranged from 0 to 3 and were derived from the product of staining intensity (0–3^+^). A zero score was considered negative, score 1 was weak positive, score 2 was moderate positive and score 3 was strong positive. (2) The percentage of positive tumours stained (on a scale of 0−3).

### Collagen content measurement in cells

as briefly described in following: MCF7 and AA-MCF7 cells were seeded on a glass bottom plate stained with CNA-35-GFP and fixed with 4% paraformaldehyde. The plates were imaged using a confocal microscope equipped with SHG system enabling us to image fluorescent and SHG at the same time. The results showed more collagen and fibrilar structure in both fluorescent imaging of CNA-35-GFP and SHG respectively (Figure 2C).

### Statistical analysis

A two-tailed unpaired Student’s *t*-test was performed to compare treated and non-treated groups of animals and also for tumours against normal cells to determine statistical significance. The significance level was set as *P*<0.05.

### Ethics statement

All procedures on animals were carried out in compliance with the Guide for the Care and Use of Laboratory Animal Resources (1996), National Research Council, and approved by the Institutional Animal Care and Use Committee, University of South Florida (IACUC# R4051)

