## Supplementary material for "Collagen Production and Niche Engineering: A Novel Strategy for Cancer Cells to Survive Acidosis and Evolve": Supplemntal materials

### Supplementary Information:

#### Figure S1.

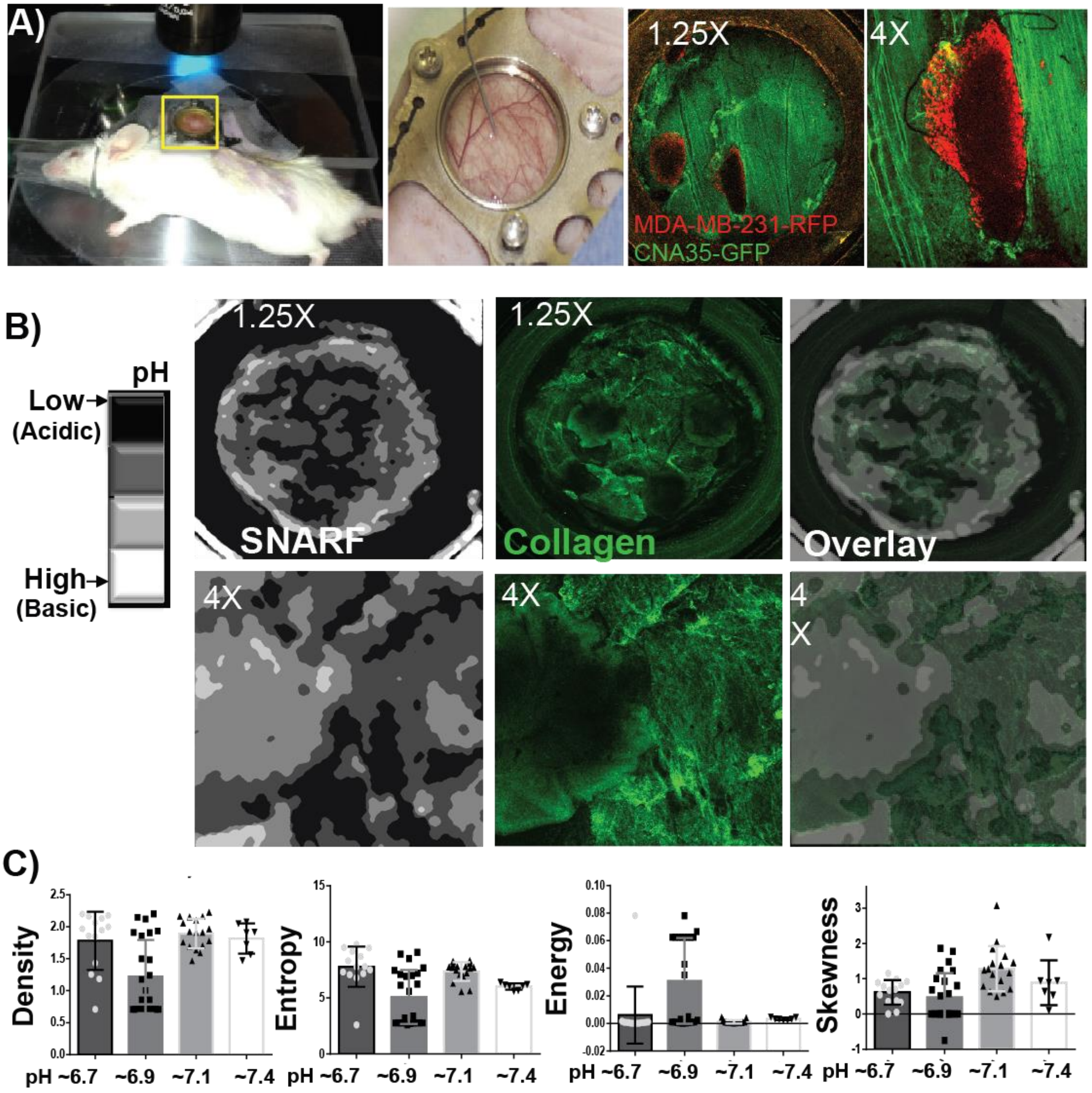


**Fig. S1. Dorsal window chamber (DWC) experimental design to study the collagen structure in different acute pH environment. A and B)** We designed a new DWC experiment by adding collagen marker directly to the window area to mark the collagen, CAN-35-GFP was added at the day of surgery and was renewed every week. The pH of whole DWC region was measured using SNARF1-AM which is a pH sensitive die and has different fluorescent emissions in different pHs. The advantage of this technique is that we are enabled to study the collagen structure and pH of the window area at the same time and real time in animal model. **C)** Our results showed that collagen structure changes is not very much related to local acute pH changes and must come from the cells.

**Figure S2.**

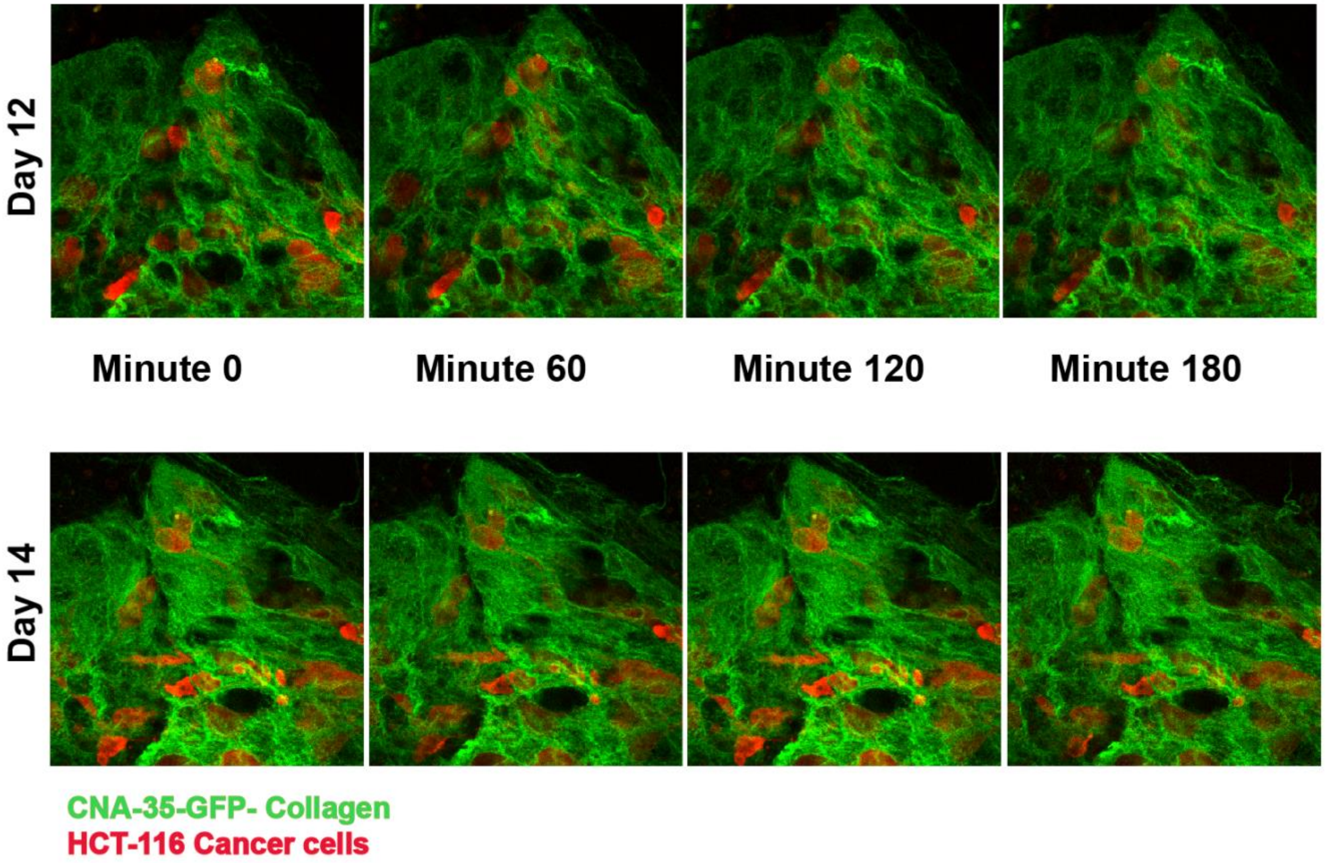


**Fig. S2. Real time dorsal window chamber (DWC) study of the collagen and cancer cells.** Our developed DWC experiment is used to capture the real time collagen structure modification by cancer cells.

**Figure S3.**

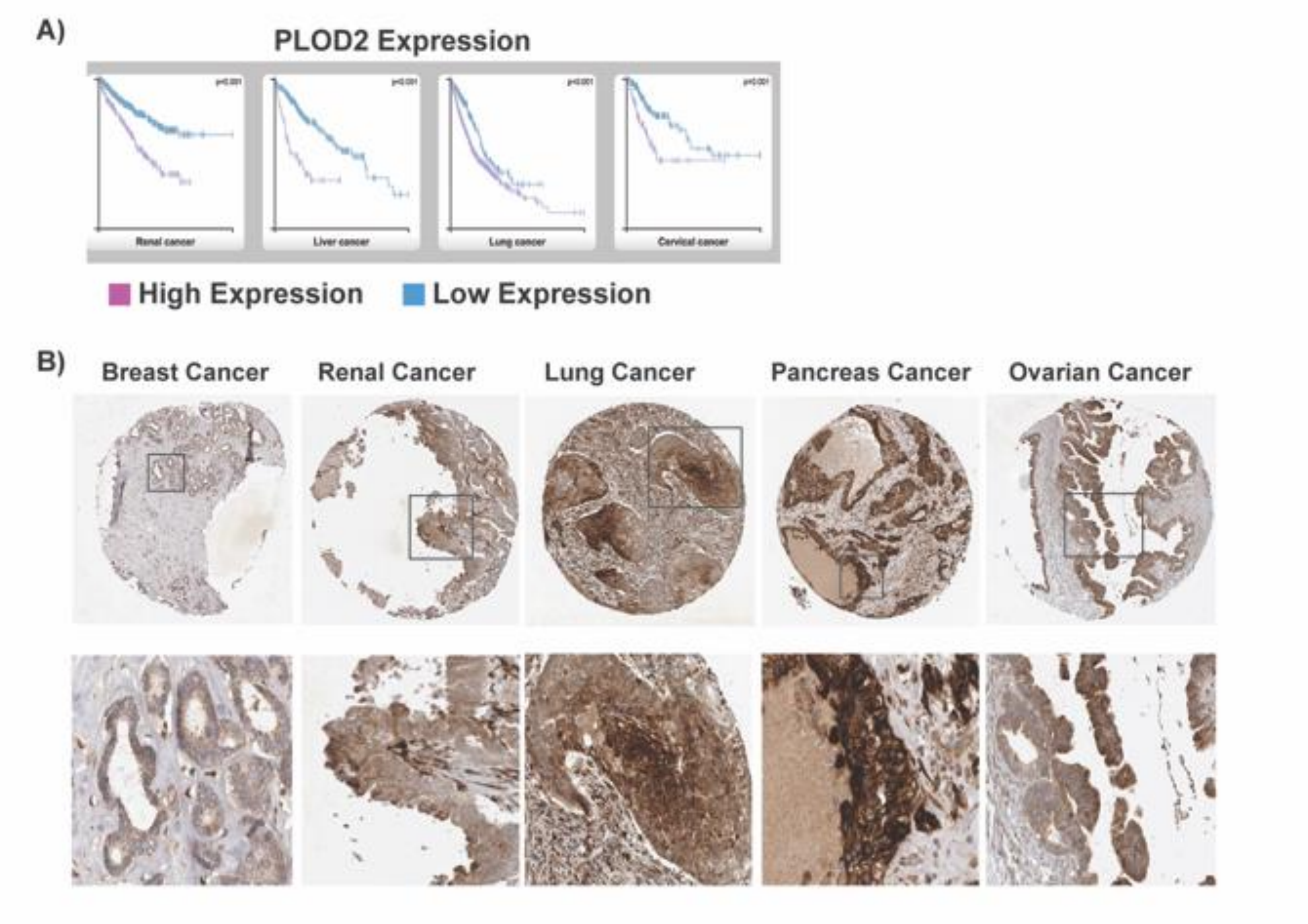


**Fig. S3. PLOD expression in different cancer type from protein atlas.** A) Overall survival of patients in different cancer type over PLOD2 expression. High PLOD2 expression significantly decreases the survival of patients. B) Representative images of biopsy of different cancer types at the early stages with high expression PLOD2. Underneath each biopsy is the zoom of selected region that implies the higher expression of PLOD2 expression at the tip of growing tumors toward the inside the duct that shows the role of collagen production in cancer cells growth inside the duct. Pictures and data are extracted form human protein atlas website: www.proteinatlas.org.

**Figure S4.**

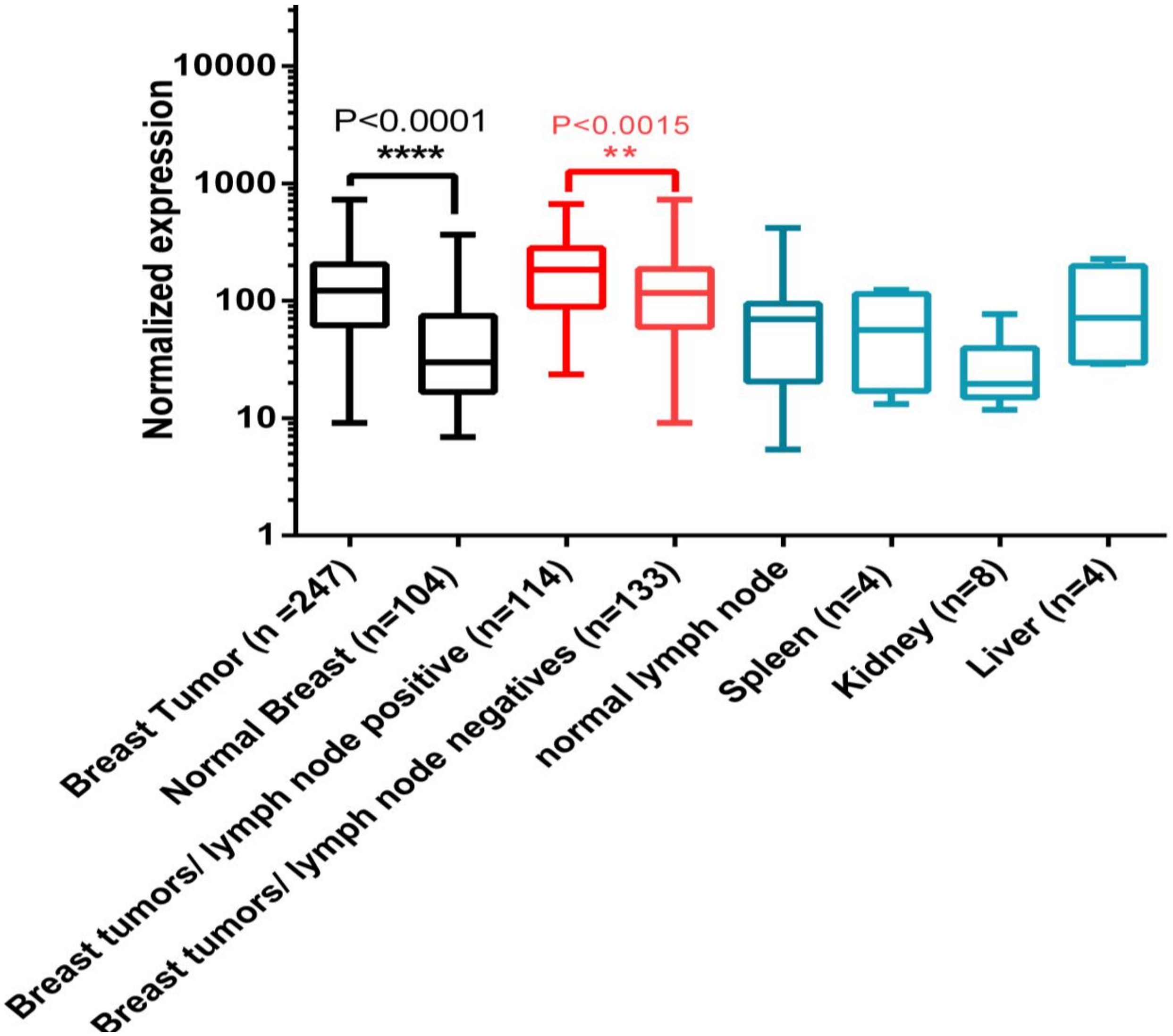


**Fig. S4. Microarray analysis of Moffitt patient's data.** Breast tumors have significantly higher amount of TGM2 compared to adjacent normal samples and other normal organs. TGM2 expression is also higher in patients with metastatic to lymph node compared to non-metastatic ones.

**Figure S5.**

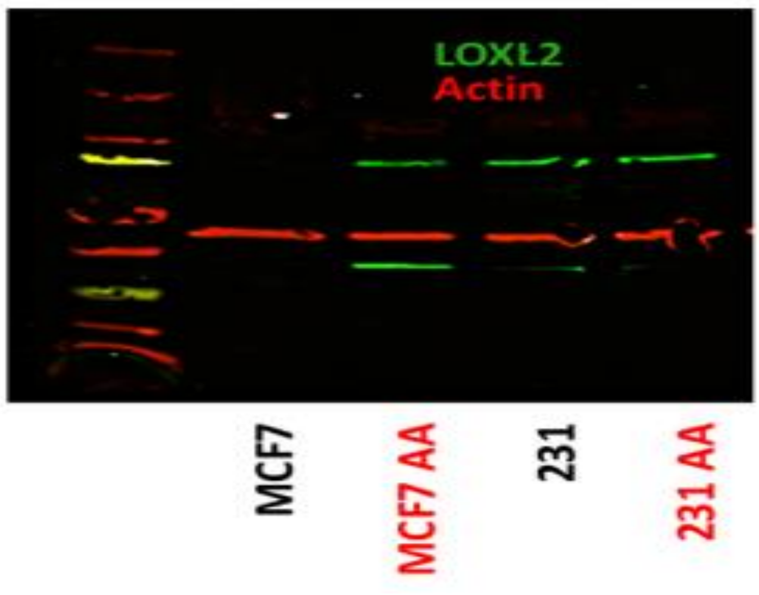


**Fig. S5. LOXL2 in acid-adapted cancer cells.** Acid adapted MCF-7 cells have higher amount of LOXL2, an enzyme that has been shown play role in collagen crosslinking and stability. Western blot approved the higher expression of LOXL2 in acid adapted MCF-7 cells vs the non-adapted one. There is no change in MDA-mb-231 cells content of LOXL2. From the secretome data we assume that they use more TGM2 than LOXL2 so they didn’t need to increase the expression.

**Figure S6.**

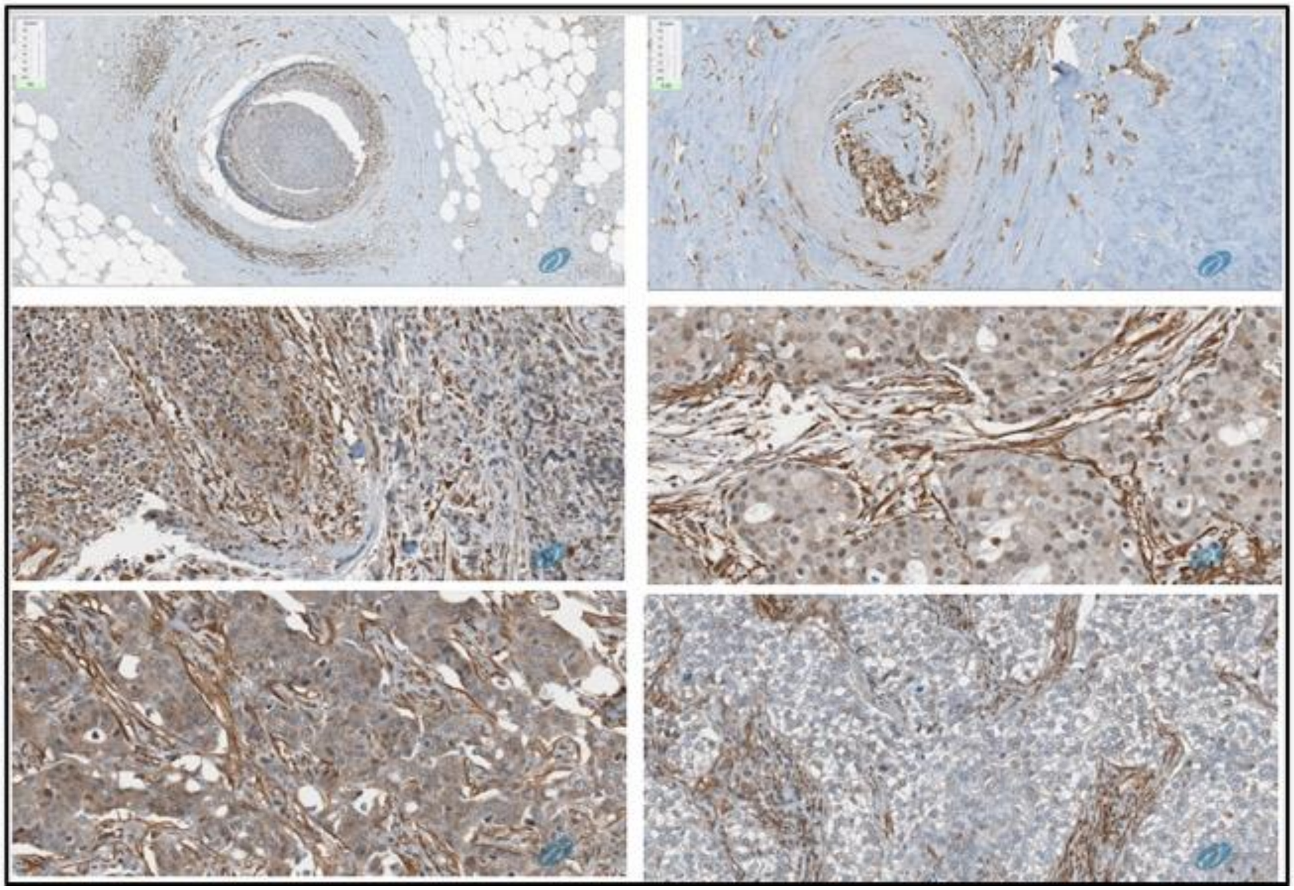

**Fig. S6. TGM2 expression in whole mount breast tumors.** We see a lot of TGM2 protein in the acidic regions such as center of DCIS (top two images) and in the extracellular spaces and interacting with fibers confirming the crosslinking role of this protein.

**Figure S7.**

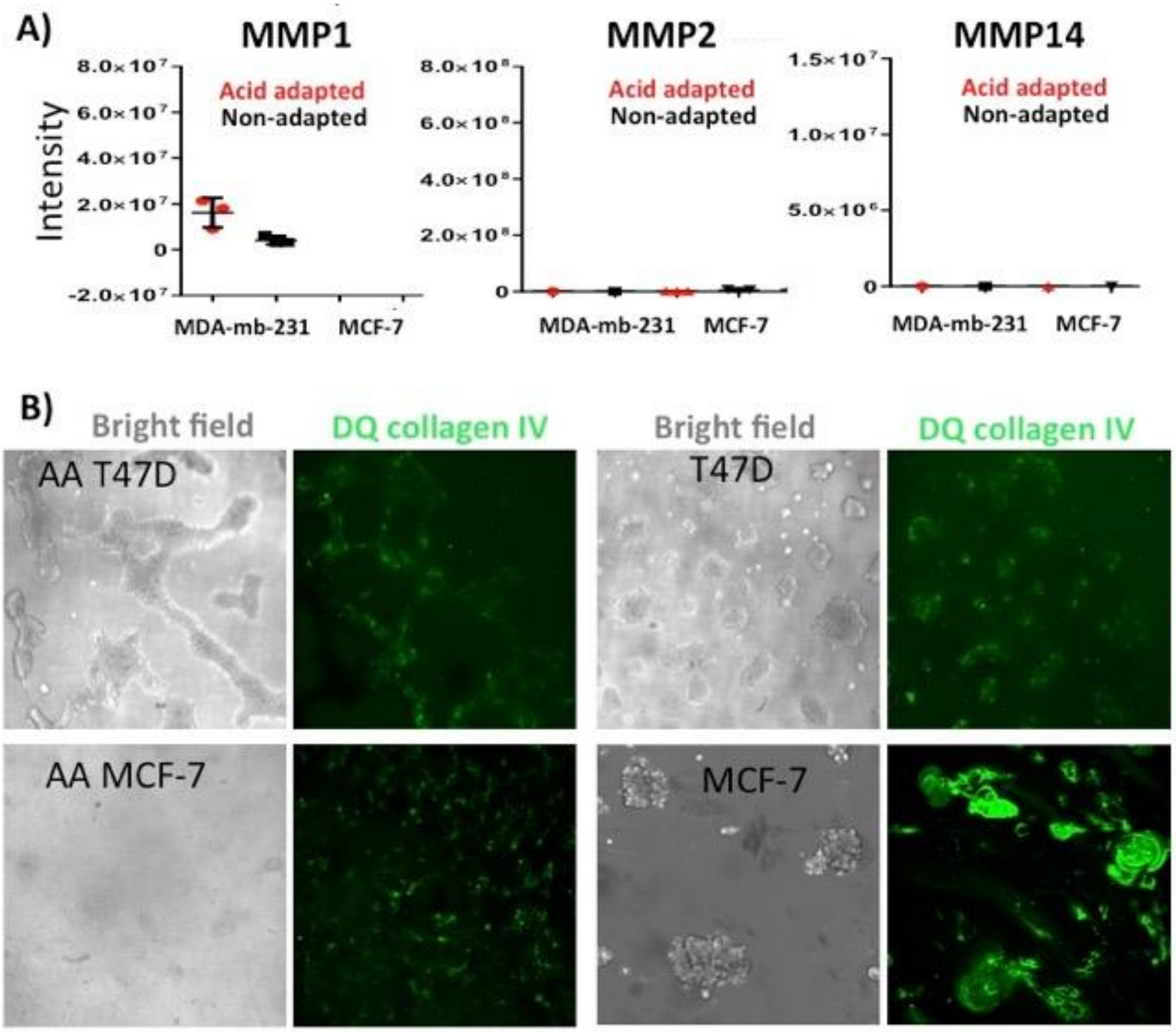


**Fig. S7. Expression statues of MMPs in acid adapted and non-adapted cells. A)** The MMPs didn't show anu change in secretome analysis. B) DQ collagen experiment was conducted to approve the secretome finding.No difference was found between acid adapted and non adapted MCF-7 and T47D cancer cells.

**Figure S8.**

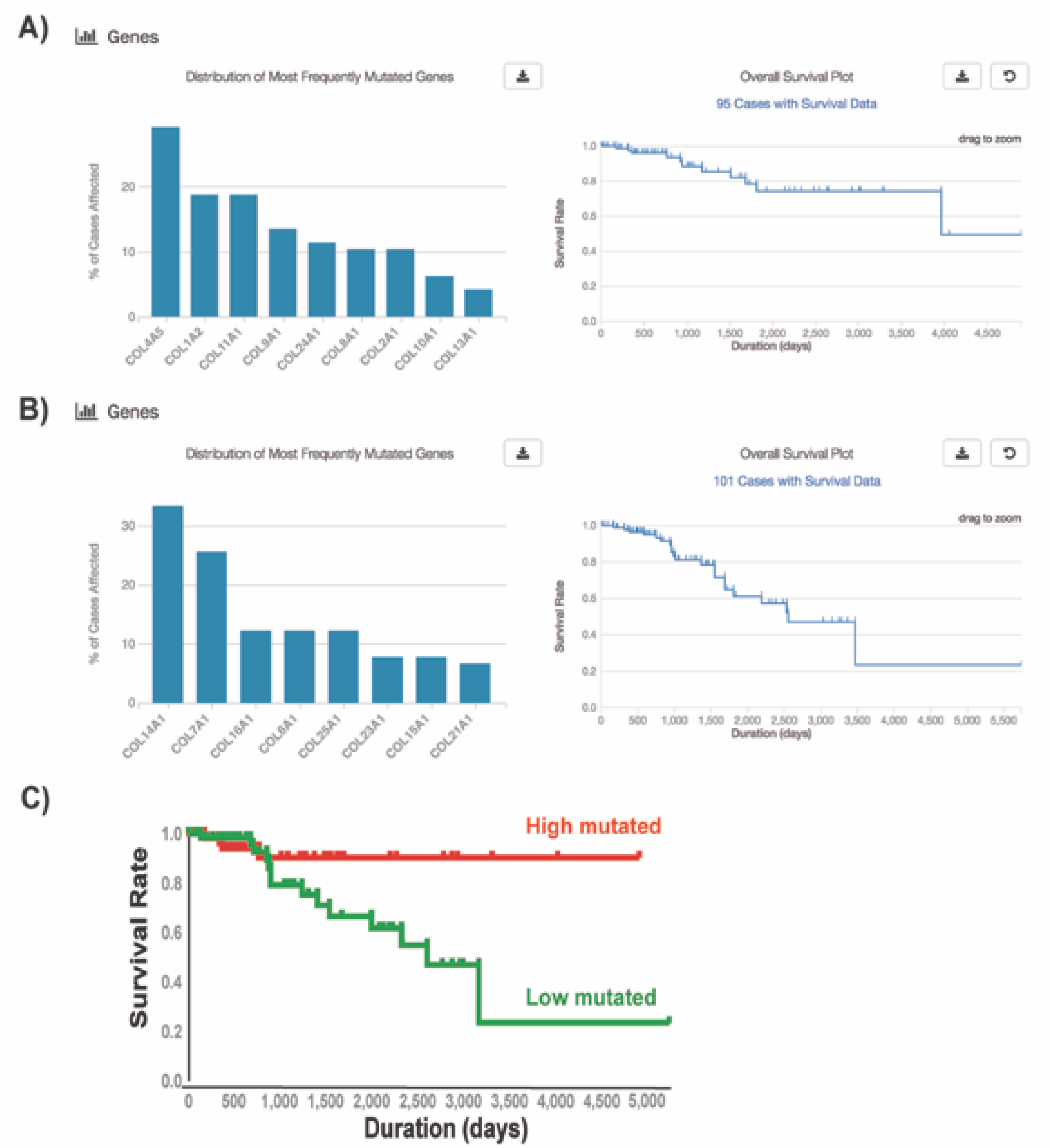


**Fig. S8.** A and B) TCGA analysis of patient overall survival with mutation in bad and good collagens. C) Survival analysis of patients with mutation in different collagen genes. Based on the expression level analysis showed in A, collagens were categorized in two groups: High expressing collagens (bad collagens) and low expressing collagens (good collagens). The overall survival of patient with high collagen expression mutations is higher than patient with mutations in low collagen genes.

Figure S9:

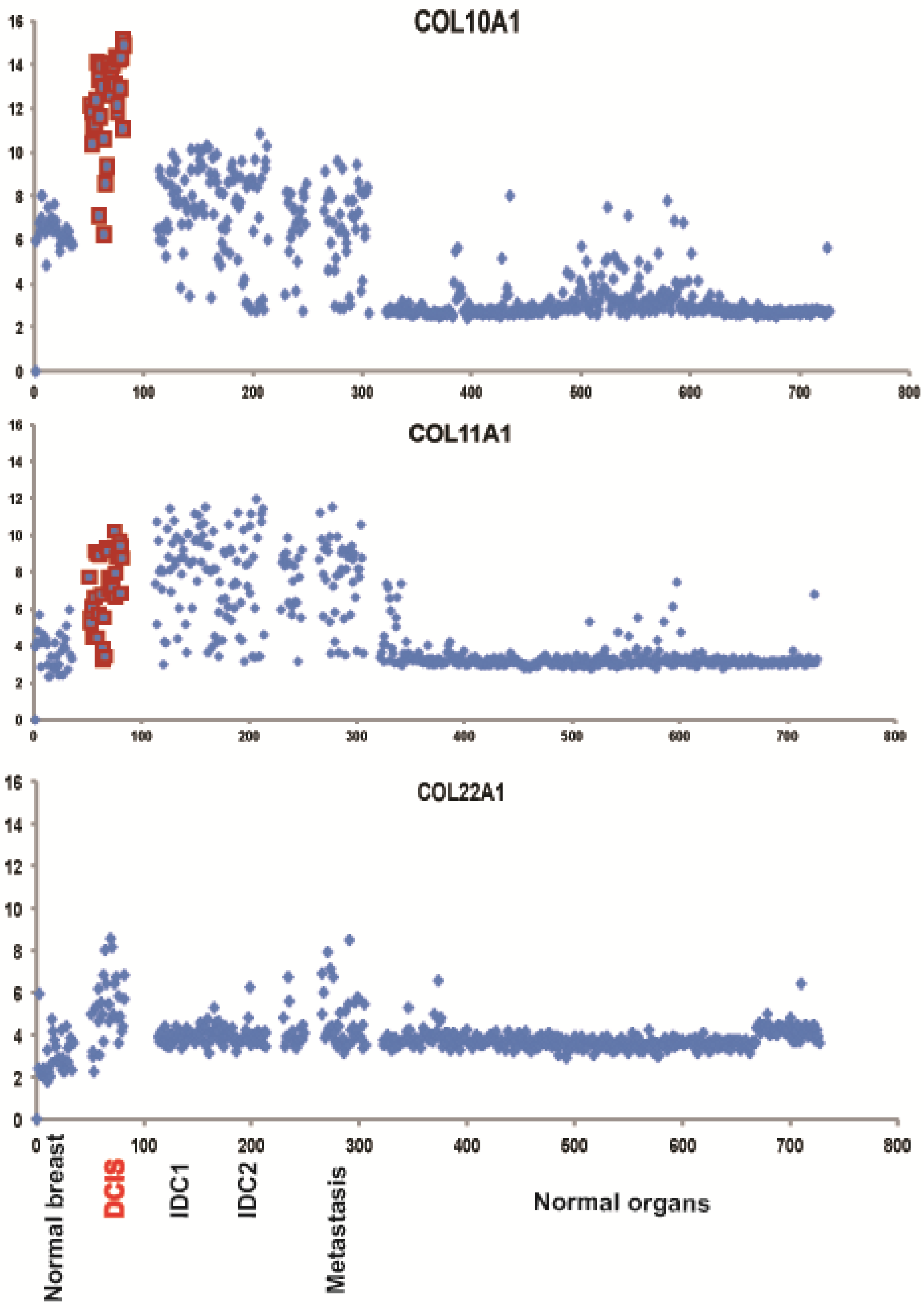


**Fig. S9.** Differential expression analysis of Moffitt Cancer Center patient's data showed correlation with TCGA data. In our database three collagens were covered: Col10a1, Col11a1, and Col22a1 that Col10a1 and Col11a1 are overexpressed in breast tumor samples while Col22a1 has no change.
